## Supporting Information for "Denoising Reveals Low-Occupancy Populations in Protein Crystals"

#### Supporting Notes

##### S1. Measuring Gaussianity for the Voxel-Value Distribution of a Difference Map

Common quantitative measures of Gaussianity for a random variable include skewness, kurtosis, and negentropy [1–3]. We construct a simple statistical model to evaluate the properties of these measures as indicators of difference map quality. In our model, we assume that individual voxels within the map are identically and independently distributed [4,5], each taking a value that represents either positive density (+1), negative density (-1), or no change (0). We incorporate Gaussian noise in the model to simulate noisy electron density.

Each voxel value, $x_i$, is sampled as follows:

$$x_i \sim \begin{cases} \mathcal{N}(0, \sigma_n^2) & \text{with probability } n \\ \mathcal{N}(-1, \sigma_n^2) & \text{with probability } \frac{1}{2}(1-n) \\ \mathcal{N}(+1, \sigma_n^2) & \text{with probability } \frac{1}{2}(1-n) \end{cases}$$

where $n$ denotes the fraction of background pixels, and $\sigma_n$ is the noise standard deviation.

We perform random sampling from these distributions over varying fractions of background ($0 \leq n \leq 1$, see Figure S1(a)). Ideally, a quality indicator for difference electron density (DED) maps would vary monotonically with respect to the amount of signal.

Because our model produces difference density histograms that are symmetric around zero, the skewness remains constantly zero (Figure S1(b)), indicating that skewness does not always reflect changes in noise and is therefore unsuitable as a noise indicator in this context. Figure S1(c-d) show kurtosis and negentropy as functions of the fraction of background: we test models with noise components of increasing standard deviations ($\sigma_n$) and find that negentropy exhibits a clear monotonic decrease with increasing background noise, consistent across all tested values of $\sigma_n$. Kurtosis, on the other hand, behaves differently, crossing zero as the background fraction rises, indicating a more complex response.

Based on these observations, we select negentropy as the preferred indicator and proceed to test whether maximizing negentropy could serve as an effective method for ranking the signal-to-noise ratio in difference density maps.

##### S2. Validation of the Choice of Regularization Parameter for Total Variation Denoising

As introduced in the main text, the choice of the $\lambda$ regularization parameter is critical in total variation (TV) denoising; it determines the level of denoising at the expense of fidelity to the original data, as formulated by the original expression for the TV minimization problem by Rudin, Osher, and Fatemi [6] and implemented by Chambolle [7]:

$$\min_u \sum_{i=0}^{N-1} \left( |\nabla u_i| + \frac{(s_i - u_i)^2}{2\lambda} \right) \tag{1}$$

where the objective is to recover the signal $u$, characterized by a lower total variation (defined as $\sum_{i=0}^{N-1} |\nabla u_i|$) than the original signal $s$, while ensuring $u$ closely resembles $s$ in an $L^2$ sense. For $\lambda$ close to zero, the output signal is almost unchanged from the original. As $\lambda$ increases, the term minimizing total variation dominates, leading to increased smoothing and decreased fidelity to the input data.

To determine an optimal regularization value for a DED map, we propose selecting the value of $\lambda$ that maximizes the negentropy of the voxel value array. We validate this approach by calculating a synthetic *trans*-to-*cis* difference density map (Figure S3(a)). To introduce noise, we add random values to the *trans* and *cis* structure factors, drawn from a Gaussian distribution with a mean of 0 and a standard deviation equal to 10% the standard deviation of the structure factor magnitudes, to resemble experimental maps. The real space map displays the signal as well as the expected added noise (Figure S3(b)).

Figure S3(c) presents the negentropy values obtained by denoising the map with a range of regularization weights. Notably, the maximum negentropy corresponds closely to the value of $\lambda$ that also maximizes the real-space Pearson correlation coefficient between the denoised map and the ground truth synthetic map density ($\rho_{\text{true}}$).

**S3. Validation of Iterative Total Variation Denoising with a Synthetic Noisy Map**

In addition to the experimental data presented in the main text, we also test the iterative-TV density modification technique (it-TV) on the synthetic noisy map described in the previous section (Figure S3(b)). it-TV produces a map that preserves the expected positive and negative densities while significantly reducing noise (Figure S3(d)). The diagram in Figure S3(d) compares the noise-free ground truth map ($\rho_{\text{true}}$) with the denoised map at each iteration ($\rho_{\text{itTV}}$). As $\rho_{\text{itTV}}$ iteratively improves, it more closely approximates $\rho_{\text{true}}$, with its negentropy converging to a maximum.

This alignment between negentropy optimization and the maximization of the Pearson correlation coefficient with $\rho_{\text{true}}$ in synthetic noisy data for both TV and it-TV treatments further supports the use of negentropy as an effective quality statistic.

### Supporting Figures

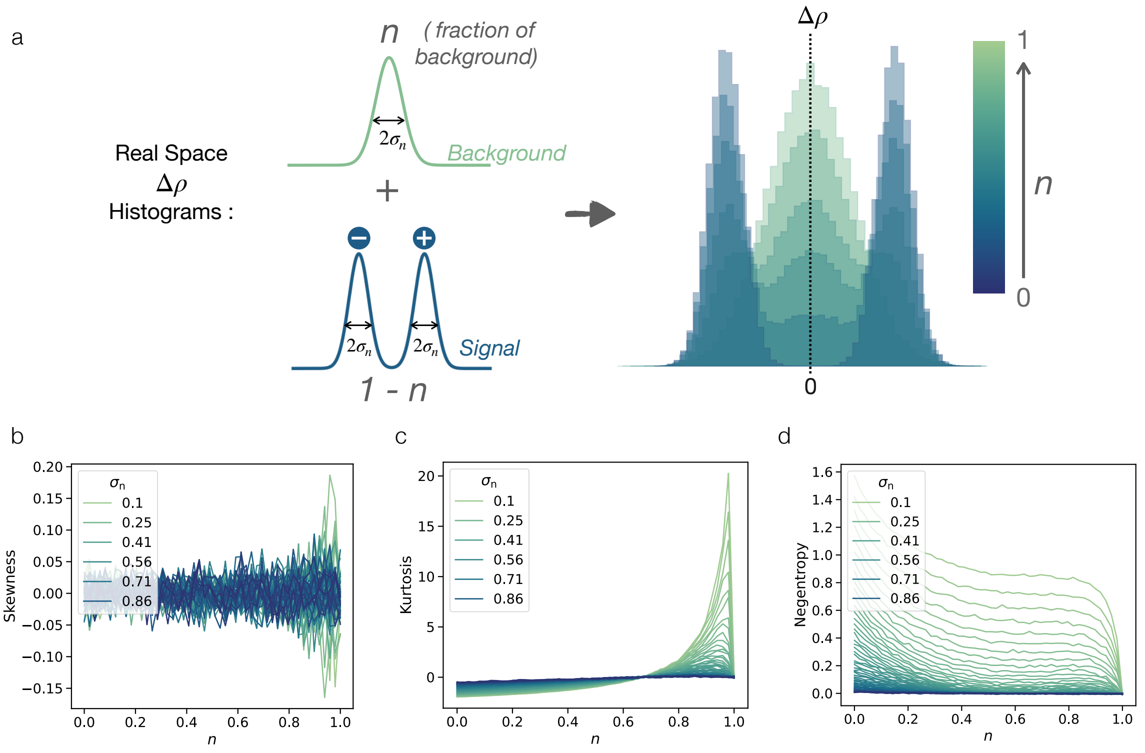


**Figure S1. Quantitative indicators of Gaussianity in simulated difference electron density (DED) maps as a function of noise fraction. (a)** A model for difference density, where voxel values are sampled from a Gaussian distribution with probabilities determined by the noise fraction $n$. Each voxel is assigned a value of positive density (+1), negative density (-1), or no change (0), with Gaussian noise added at standard deviation $\sigma_n$. **(b-d)** Skewness, kurtosis, and negentropy measure deviations from normality [1–3]. The plots show these metrics as the noise standard deviation in the model varies ($0 \leq \sigma_n \leq 1$). The results indicate that negentropy, which decreases monotonically with noise, is a reliable indicator of noise levels in DED maps. On the other hand, skewness remains zero, as expected for symmetric histograms generated by this model, and kurtosis behaves non-monotonically, crossing zero as a function of noise fraction.

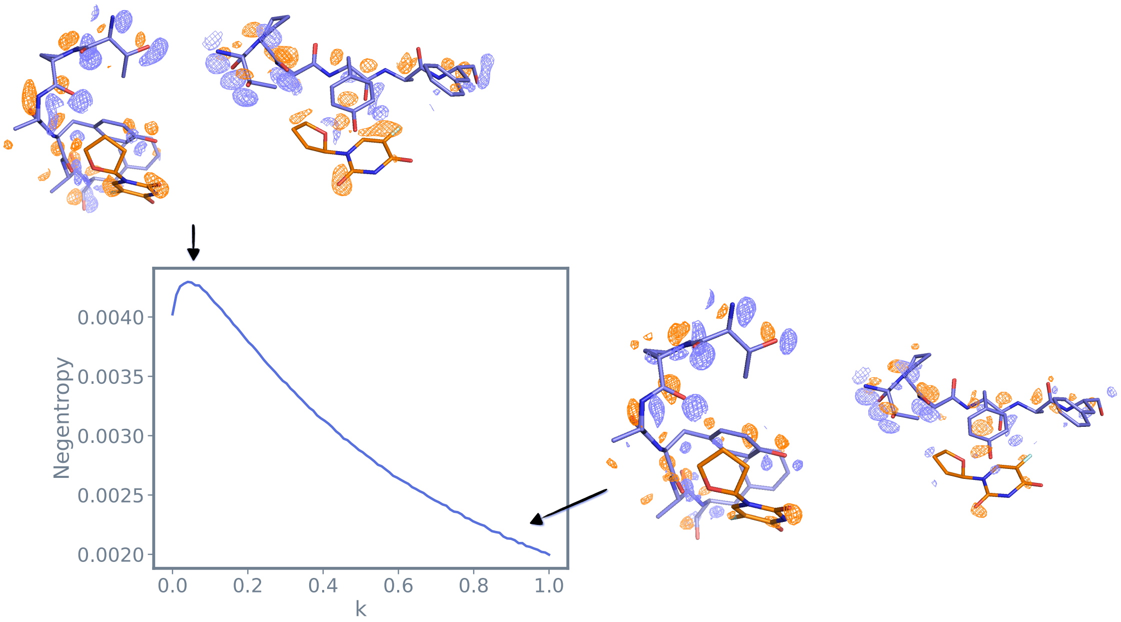


**Figure S2. Negentropy can be used for parameter determination in k-weighting of difference structure factors**. The choice of the *k* parameter for outlier rejection in k-weighting [8,9] has so far been determined by user visual inspection. We show here that negentropy maximization can be used to choose the *k* value for the $M^{pro}$-tegafur data (PDB ID 7AWR) .

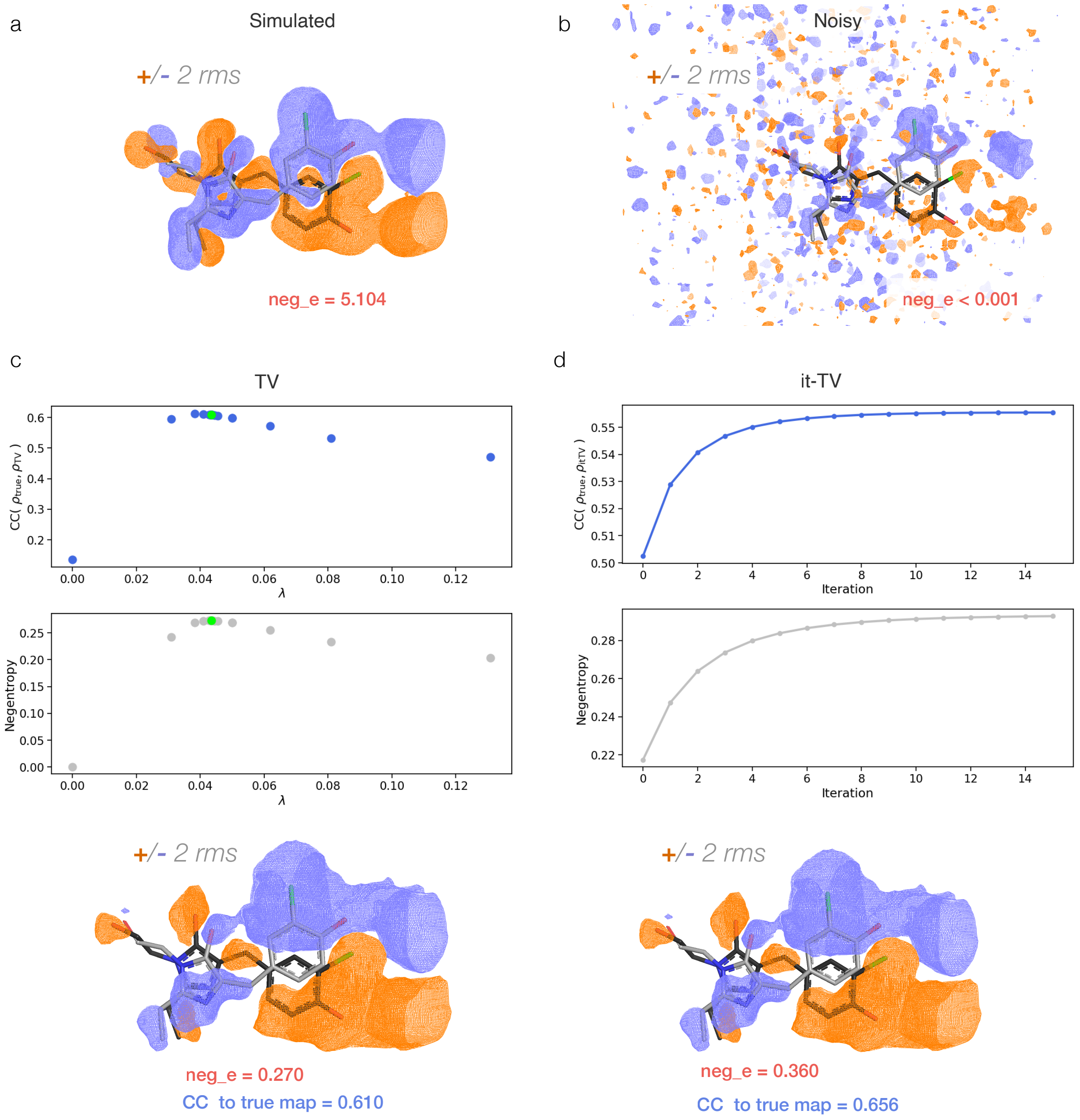


**Figure S3. Validation of total variation denoising with a synthetic noisy map.** **(a)** Simulated *trans*-to-*cis* difference density map. **(b)** Noise introduced by adding random values to the *trans* and *cis* structure factors, sampled from a Gaussian distribution with mean 0 and standard deviation equal to 10% the standard deviation of the structure factor magnitudes. The real-space map thus displays both the signal and the added noise. **(c)** Negentropy values for the denoised map across a range of regularization weights. The peak negentropy closely matches the $\lambda$ value that maximizes the real-space Pearson correlation coefficient between the denoised map and the true calculated map ($\rho_{\mathrm{true}}$). **(d)** Application of the iterative-TV (it-TV) technique further refines the denoised map and results in improved negentropy and correlation to ground truth map values. As iterations progress, the denoised map ($\rho_{\mathrm{itTV}}$) increasingly approximates the ground truth map ($\rho_{\mathrm{true}}$), with negentropy plateauing to a maximum, supporting its use as a robust quality metric.

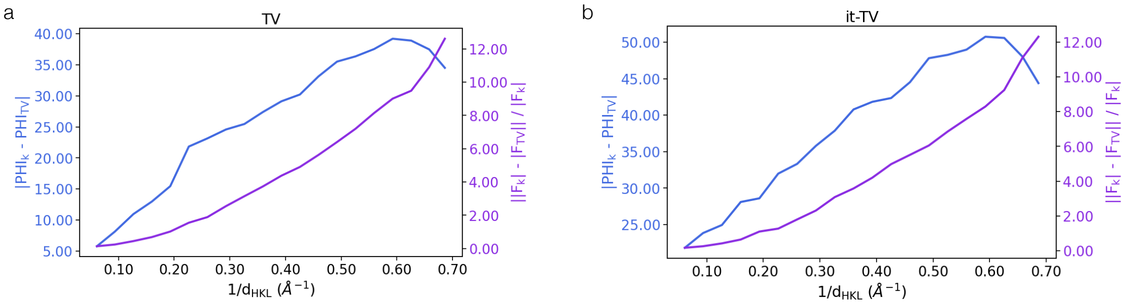


**Figure S4. TV denoising primarily modifies high resolution reflections.** Structure factor amplitudes and phase differences between the k-weighted map for the PDB ID 8A6G Cl-rsEGFP2 test case and the map generated from **(a)** single pass TV denoising and **(b)** iterative-TV. The average statistics are plotted after binning the datasets in 20 resolution shells.

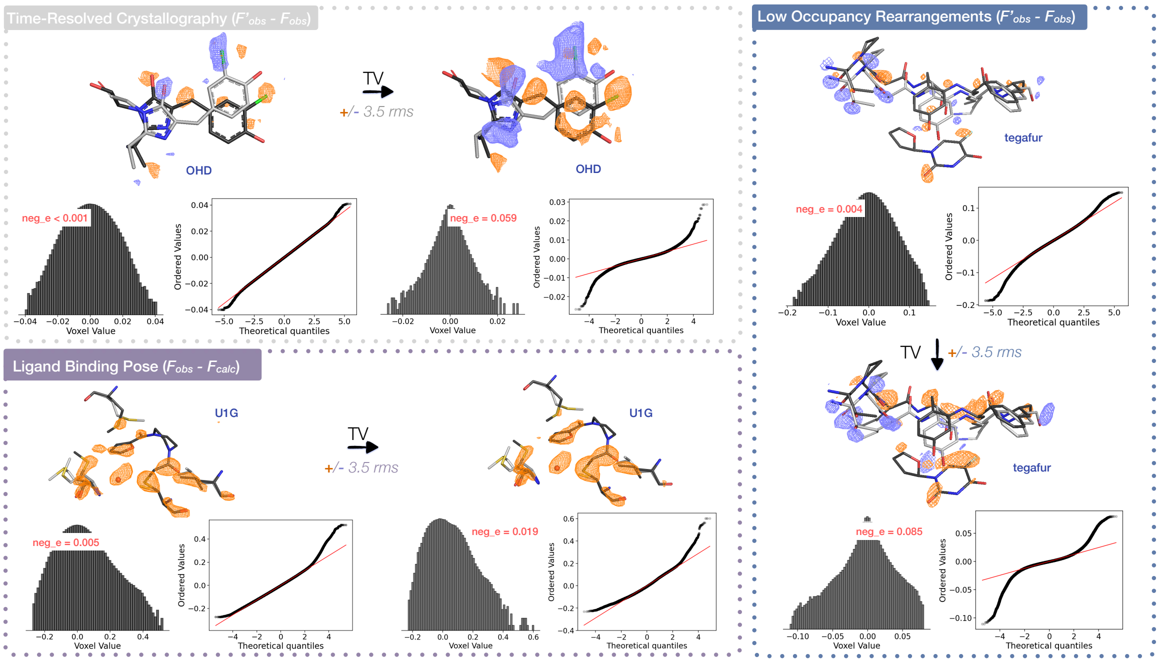


**Figure S5. Map denoising through TV minimization is applicable to a range of science cases.** The best negentropy DED map after a single pass of TV denoising at its refined $\lambda$ is displayed for the three crystallographic case studies outlined in the text. Reference state structures are shown in gray, while structures that were refined to the perturbed dataset are shown in black. Voxel value histograms for the maps (with a log-scale on the y-axis), probability plots, and respective negentropy values (neg_e) are also reported.

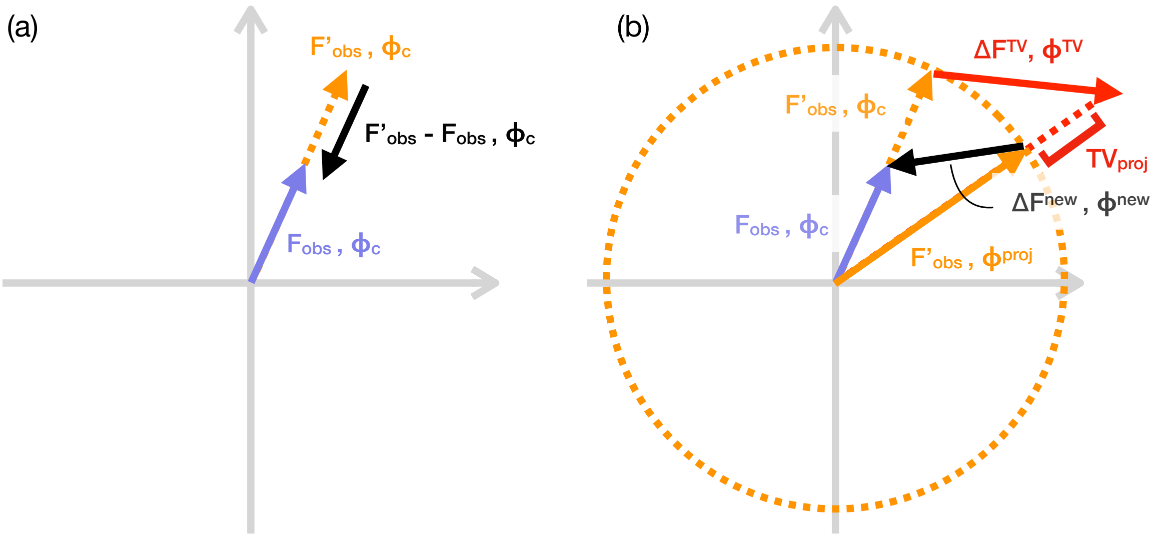


**Figure S6. TV-denoised structure factors can be used to better estimate the phases of an F' state. (a)** Fourier difference syntheses are based on the assumption of isomorphism between the native **F** and the derivative **F'**. The approximation of using the phases from the reference state ($\phi_c$) for both **F** and **F'** introduces a source of error, which is expected to halve the signal-to-noise ratio in the final map [10] **(b)** We postulate that projecting the vector of TV-denoised structure factors, $\Delta\mathbf{F}^{\mathbf{TV}}$, onto the phase circle with magnitude $|F'_{obs}|$, can provide an improved estimate for the **F'** phases, i.e. $\phi^{\text{proj}}$. A better estimate for the difference structure factors can therefore be computed: $\Delta$**Fs** $\approx \Delta\text{F}^{\text{new}}$, $\phi^{\text{new}}$.

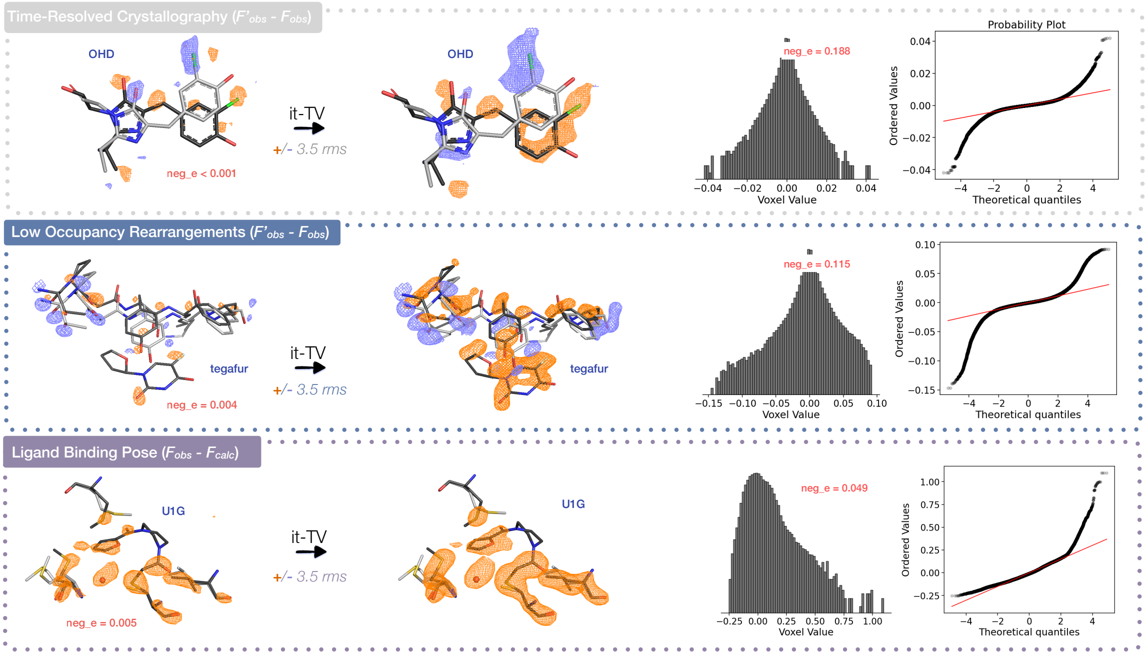


**Figure S7. An iterative TV minimization algorithm estimates the phases for low occupancy states.** We show it-TV maps for our three test datasets, with their voxel value histogram, probability plots, and associated negentropy. Reference state structures are shown in gray, while structures that were refined to the perturbed dataset are shown in black.

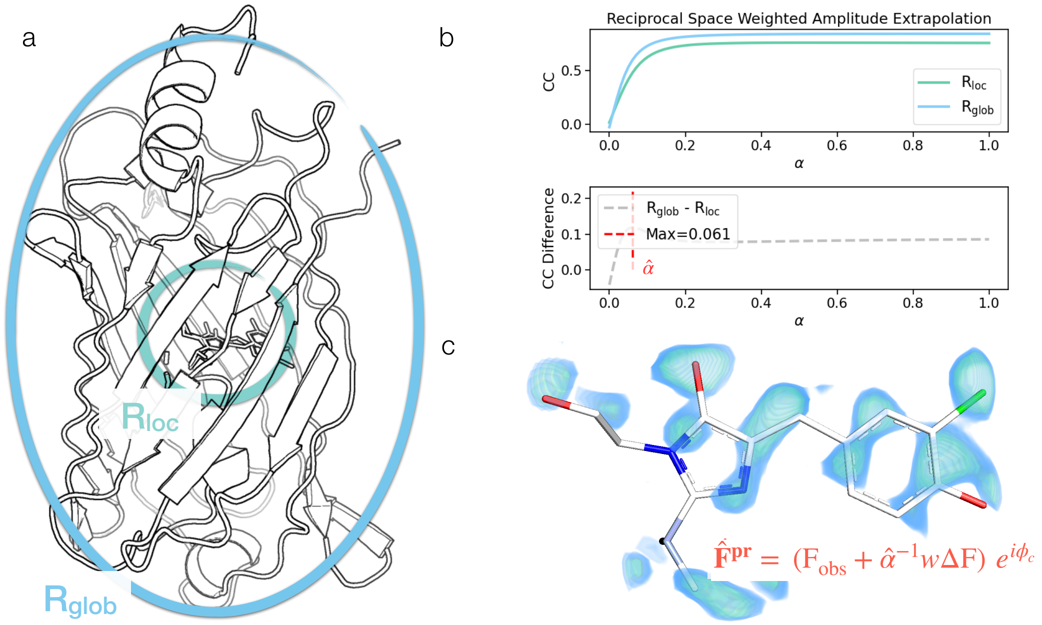


**Figure S8. Reciprocal space background subtraction for extrapolating the density of perturbed states. (a)** To find an estimate for the extrapolation factor $\alpha$, we re-implement the background correction carried out by the PanDDA suite [11]: in real space, we define a local region, $\mathrm{R_{loc}}$, based on where the strongest signals in the METEOR it-TV map are found (for the Cl-rsEGFP2 chromophore pocket shown here, $\mathrm{R_{loc}}$ is set as a 5Å sphere centered on the chromophore isomerizing bond). The entire protein is defined as $\mathrm{R_{glob}}$ after a solvent mask is applied. Fourier amplitudes of the form: $\mathrm{F^{pr}} = (\mathrm{F_{obs}} + \alpha^{-1} w \Delta \mathrm{F})$ are computed for a range $0 \leq \alpha \leq 1$. The phases from the reference model ($\phi_c$) are used for map generation. **(b)** For each value of $\alpha$, the Pearson correlation coefficient between the respective $\mathbf{F^{pr}}$ map and the map obtained from the reference model calculated structure factors ($\mathrm{F_c}$) is computed. This is done for both $\mathrm{R_{loc}}$ and $\mathrm{R_{glob}}$. The fraction of $\alpha$ that maximizes the difference between these two correlation coefficients ($\hat{\alpha}$) is chosen to approximate the structure factors of the perturbed state $\mathbf{F^{pr}}$. **(c)** The extrapolated map obtained for the picosecond Cl-rsEGFP2 photoisomerization data is shown: the outline of the *cis* photoproduct species becomes clear and can confidently support the presence of the new species.

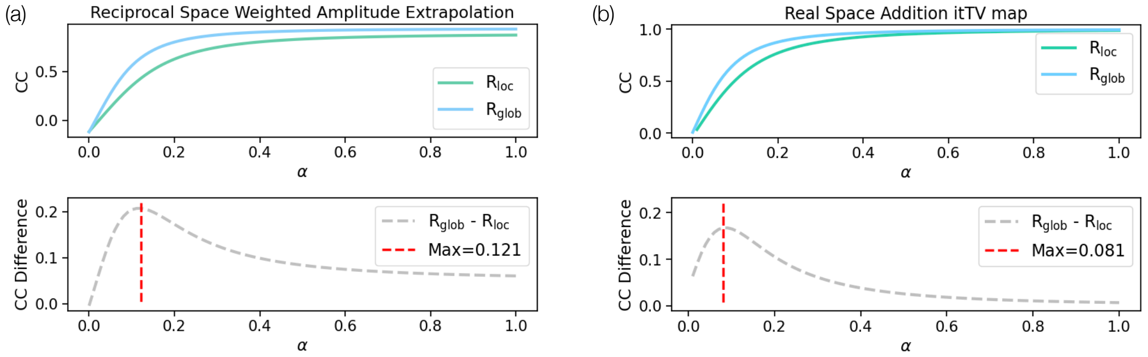


**Figure S9. Estimate of $\alpha$ parameter for map extrapolation for the $\mathrm{M^{pro}}$-tegafur complex.** Determination of the $\alpha$ parameter for map extrapolation is carried out with our reciprocal space **(a)** and real space **(b)** implementations as described in the Methods section. For both cases, the local sphere is centered on the ligand and chosen with a radius of 8 Å. Note that, when the it-TV map is used for the extrapolation, the expectation that $\alpha$ should be half of the perturbed state occupancy no longer holds (as we are no longer approximating **F** and **F'** using the same phase).

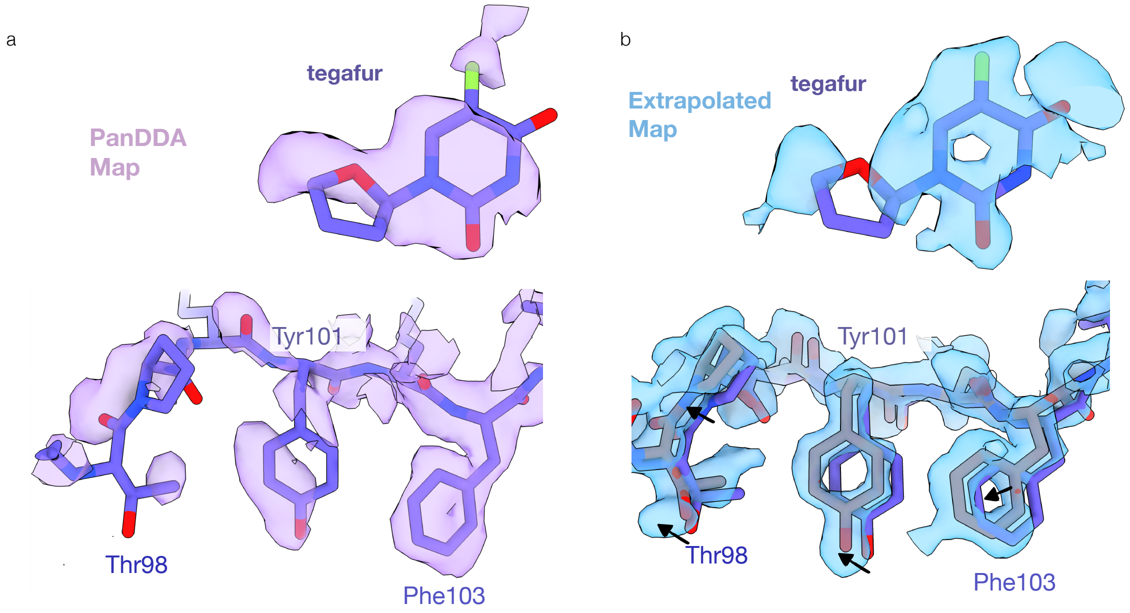


**Figure S10. Extrapolated density for the M$^{\text{pro}}$-tegafur complex obtained using reciprocal space background subtraction** The PanDDA map **(a)** for the M$^{\text{pro}}$-tegafur dataset (PDB ID 7AWR) is compared to the $\hat{F}^{pr}$ map **(b)** obtained from the reciprocal space map extrapolation procedure outlined in Figures S8 and S9(a). The extrapolated $\hat{F}^{pr}$ map displays clear outlines for Thr98, Tyr101, and Phe103 as well as backbone density that can be interpreted as rearrangements away from the fragment binding pocket and that can be used for refinement of new atomic positions. The resolution in the $\hat{F}^{pr}$ map is improved compared to the PanDDA map: ring structures and oxygen densities are considerably more interpretable for both the fragment density and the protein side chains.

a

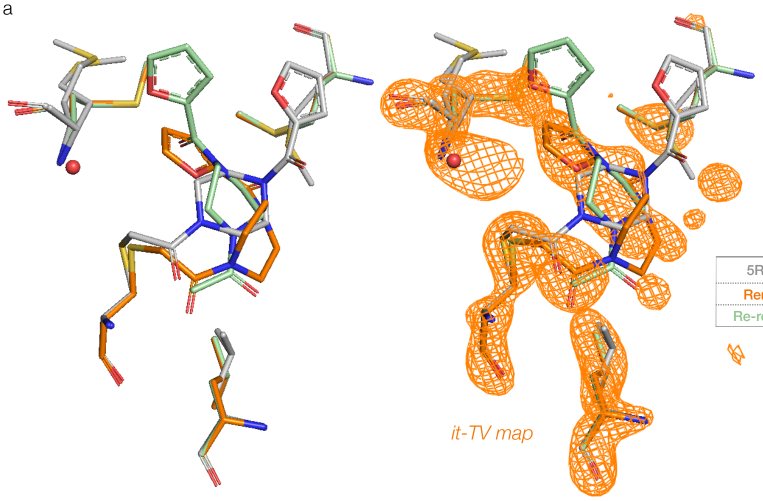


b

| | Ligand RSCC | R-work | R-free |
|---|---|---|---|
| 5RGO – deposited | 0.513 | 0.1952 | 0.234 |
| Remodeled to it-TV | **0.630** | **0.1716** | **0.230** |
| Re-refined with phenix | 0.487 | 0.1717 | 0.227 |

**Figure S11. Ligand binding pose modeling to iterative-TV density. (a)** The deposited structure for the M$^{pro}$-U1G complex (PDB ID 5RGO) is shown in gray. The structure that was manually remodeled to the density from the iterative-TV map derived in Figure 4 of the main text is shown in orange. The model that can be obtained by re-refining the deposited model with phenix.refine [12] is shown in green. The three structures are displayed with (right) and without (left) overlaying the it-TV map. **(b)** Real-space correlation coefficient (RSCC) values for ligand density between the model calculated map and the ligand polder map [13] (calculated from the original model and the deposited structure factors) are reported in the first column. R-factor values for the newly refined models and the original deposition are also reported. The structure that was remodeled to the it-TV map displays improved fit to the data as assessed by RSCC and R-factors.
